# Radiopharmaceutical therapy for metastatic prostate cancer: Insights from mechanistic modeling and *in silico* trials

**DOI:** 10.64898/2025.12.17.694859

**Authors:** Matteo Italia, Silvia Bordel-Vozmediano, José García Otero, Gabriel F. Calvo, Víctor M. Pérez-García

## Abstract

Radiopharmaceutical therapy (RPT) has rapidly evolved into a key precision-oncology modality, with radioligands now approved or in late-stage development for multiple solid tumors, including neuroendocrine and prostate cancers. RPT with [^177^Lu]Lu-PSMA (Prostate-Specific Membrane Antigen) has recently been approved as a life-prolonging treatment for metastatic castration-resistant prostate cancer (mCRPC), but its clinical use still relies on non-personalized, empirically chosen fixed schedules. Here, we develop a mechanistic, patient-personalizable mathematical model simulating mCRPC response and organ-at-risk toxicity during [^177^Lu]Lu-PSMA RPT. The model integrates tumor growth dynamics, radiobiological response, and organ-resolved pharmacokinetics inferred from mass data and standardized uptake values obtained from positron emission tomography studies. Parameters were derived from the literature, although the framework allows personalization by fitting to patient-specific data such as imaging and prostate-specific antigen levels. Using virtual patient (VP) cohorts generated via stochastic parameter sampling, we conducted *in silico* trials and validated the model by comparing simulated outcomes with published dosimetry and survival data for [^177^Lu]Lu-PSMA trials. We then explored dosing and scheduling strategies to optimize efficacy-toxicity trade-offs. Consolidated regimens with fewer, higher-activity injections improved overall survival (OS) *in silico* but increased toxicity, especially in kidneys. Cycle length had a weaker influence on OS within a 2–9 week window, while it clearly affected toxicity, whereas excessive delays (*>* 12 weeks) markedly reduced efficacy. Global sensitivity analysis identified tumor growth, uptake, and radiosensitivity parameters as key drivers of interpatient variability, and convergence testing confirmed robustness with respect to VP cohort size. These methodological findings illustrate how mechanistic modeling and *in silico* trials can inform the design and personalization of RPT regimens.

**Author summary:** Standard radiopharmaceutical therapy regimens for metastatic castration-resistant prostate cancer deliver the same doses at fixed intervals, without accounting for interpatient variability in tumor growth, drug uptake, or organ tolerance. Here, we focus on [^177^Lu]Lu-PSMA radiopharmaceutical therapy, which is now approved for these patients but still administered using one-size-fits-all protocols. We present a mechanistic mathematical model that simulates tumor and radiopharmaceutical dynamics in individual patients using a virtual patient framework. With appropriate patient-specific data, such as quantitative imaging and prostate-specific antigen levels, this model can be used to generate digital twins and evaluate personalized treatment strategies. By adjusting injection schedules and cycle timing in silico, we explored how standard treatment protocols could be optimized to improve survival while maintaining acceptable toxicity. We found that a 9 week treatment cycle achieved survival outcomes comparable to the standard 6 week protocol, with a significant reduction in toxicity, whereas longer cycle extensions led to loss of therapeutic efficacy. These results provide a quantitative basis for optimizing radiopharmaceutical therapy and highlight the potential of virtual patients and in silico trials to support patient-adapted treatments in modern oncology.

## 1 Introduction

Radiopharmaceutical therapy (RPT), also referred to as targeted radionuclide therapy (TRT), is a powerful and rapidly evolving modality in precision oncology, particularly within the emerging field of radiotheranostics, which uniquely combines diagnostic imaging and therapeutic interventions [1, 2]. The diagnostic component relies on quantitative Positron Emission Tomography (PET) or Single Photon Emission Computed Tomography (SPECT) imaging with radioisotopes to non-invasively evaluate the number and distribution of lesions, thereby assisting in determining patient eligibility for RPT [3]. By employing radiolabeled compounds that selectively bind to tumor-specific targets, RPT delivers targeted irradiation to cancer cells while sparing surrounding healthy tissue.

Recent advances have led to regulatory approval of modern RPTs for patients with metastatic castration-resistant prostate cancer (mCRPC) [4] and neuroendocrine tumors [5], through the use of prostate-specific membrane antigen (PSMA) and somatostatin receptor 2 (SSTR2) as molecular targets. These approvals followed successful clinical trials of [^177^Lu]Lu-PSMA-617 [6] and [^177^Lu]Lu-DOTATATE [7]. Meanwhile, numerous ongoing studies are expanding the spectrum of RPTs to address a wider range of disease targets, demonstrating strong *in vivo* performance and underscoring the promising future of theranostic-based RPT in advancing precision oncology [8].

One of the central challenges in oncology and RPT is the inter-patient heterogeneity of response [9]. Standard administration of fixed activities can lead to under or overdosing due to variability in tumor burden, pharmacokinetics, and organ function [10]. To fully realize the potential of RPT, there is a growing interest in personalizing treatment–selecting the right molecule, dosage, injection sites, timing, and combinations–tailored to each patient’s biological characteristics [3, 9].

Mathematics can play a transformative role in advancing personalized RPT [11]. Mechanistic models enable the simulation of radiopharmaceutical transport, tumor growth, radiation damage, protocol planning, and therapy response, thereby supporting virtual treatment testing and *in silico* optimization. Such approaches can reduce costs and risks for patients and shorten development times compared to traditional clinical trials [12]. Moreover, these models facilitate the generation of virtual patients (VPs) and digital twins [11], providing a safe framework for designing adaptive, optimal, and personalized therapies. By integrating mathematical modeling with imaging and patient-specific data, new opportunities emerge for personalized dosimetry, trial design, and treatment planning [13, 14].

The current standard of care for mCRPC involves six cycles of [^177^Lu]Lu-PSMA-617 administered at six-week intervals. Each cycle delivers a fixed activity of 7.4 GBq, regardless of individual tumor characteristics, resulting in a total administered activity of 44.4 GBq over approximately nine months [6]. In some patients, this regimen was effective, with minimal toxicity even among those heavily pretreated [15, 16].

Nonetheless, little is known about the potential therapeutic benefits of modifying this protocol or adapting it to patient-specific characteristics.

To address this concern, Zaid *et al*. developed a mathematical compartmental model and investigated optimal dosing [17]. Their model described activities in tumor and organs at risk (OARs), governing tumor response and OAR toxicity, respectively. By simulating different treatment schedules, the authors identified protocols that could yield higher efficacy than the current 6-cycle/6-week protocol [17].

In this study, we improved the model of Zaid *et al*. [17] by extending its biological modeling and incorporating patient-specific parameters. We validated the model by reproducing *in silico* toxicity [18, 19] and survival [20, 21] outcomes observed in different clinical trials. We further investigated the virtual cohort size required to ensure population-level reliability and avoid cohort-specific artifacts. Finally, we applied the validated model to simulate RPT delivery and therapeutic response in mCRPC, using virtual cohorts to explore the impact of patient-specific heterogeneity and treatment scheduling. Specifically, we tested regimen variations by modifying cycle dosages or cycle lengths, while keeping the total administered activity fixed at 44.4 GBq [6].

## 2 Materials and methods

### 2.1 Conceptual framework

The biological dynamics of tumor growth, radiobiological response, pharmacokinetics, and absorbed doses (ADs) in OARs in [^177^Lu]Lu-PSMA therapy considered in this paper are summarized in Fig 1. Our conceptual scheme incorporates a growing mCRPC tumor mass subject to RPT-induced cell death. Tumor cell subpopulations included in the total tumor mass (*M*_T_) are: proliferating tumor cells (*M*_TP_), cells with sublethal radiation-induced damage (*M*_TSD_), and lethally damaged cells (*M*_TD_).

**Fig 1.**
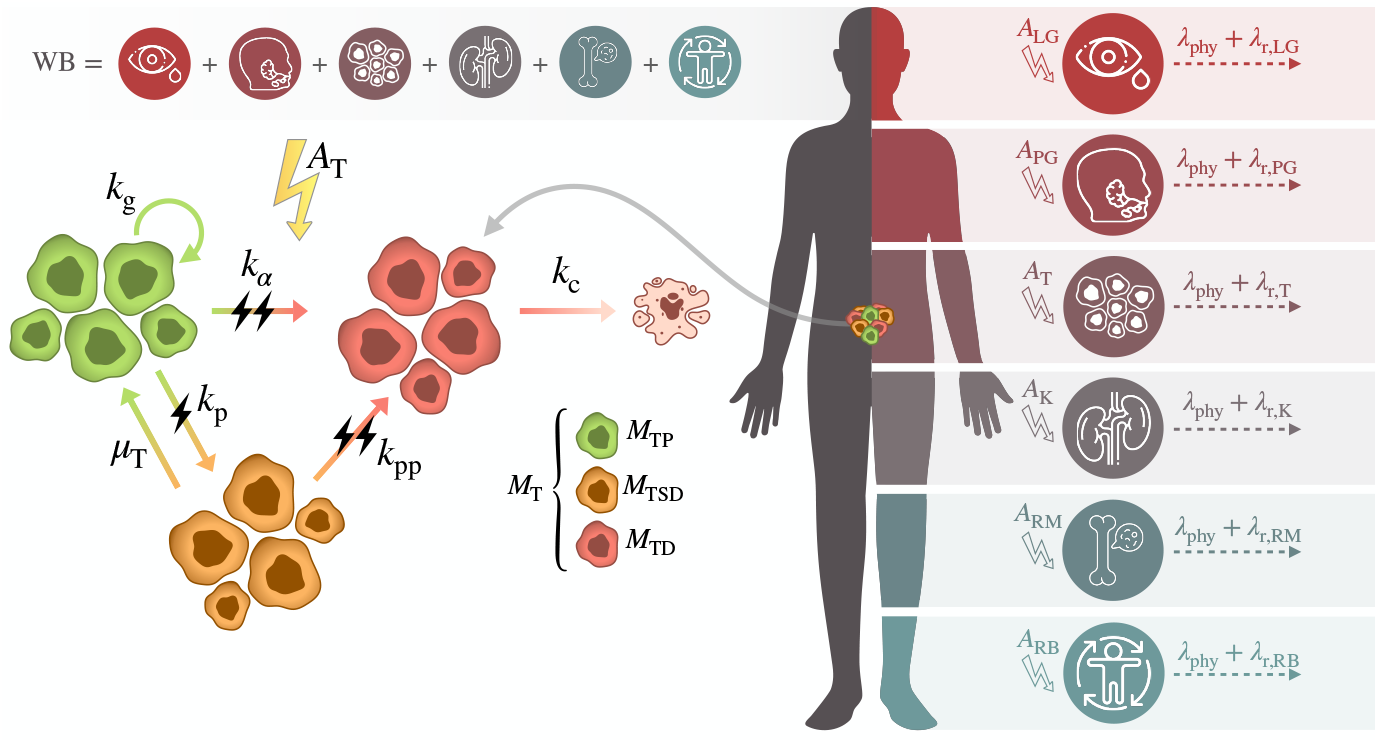
Modeling scenario for tumor-mass dynamics and organ dosimetry in radiopharmaceutical therapies. Left panel: The total tumor mass (*M*_T_) is partitioned into proliferating tumor cells (*M*_TP_), sublethally damaged cells (*M*_TSD_), and lethally damaged cells (*M*_TD_). Transitions between these states are driven by the time-varying tumor activity (*A*_T_). Right panel: The whole-body (WB) is partitioned into organs at risk (OARs), tumor lesions (T), and remaining body (RB). The included OARs are the kidneys (K), red marrow (RM), parotid glands (PG) and lacrimal glands (LG). The activity in each body compartment (*A*_X_) follows an exponential decay law governed by the physical decay rate of the radionuclide (*λ*_phy_) and the compartment-specific biological clearance rate (*λ*_r,*X*_ ).

Proliferating cells are assumed to grow following an exponential law. This behavior has been observed in a retrospective analysis of prostate-specific antigen trajectories from eight randomized clinical trials in mCRPC that used a linear combination of exponentials to describe tumor growth over time [22]. Proliferating cells can be damaged by RPT and move to the sublethal or lethal compartments. Sublethally damaged cells may repair and return to the proliferative compartment or accumulate further damage and transition to the lethally damaged compartment. Lethally damaged cells are cleared from the system with a characteristic clearance time.

OARs considered in the model were the kidneys (K), red marrow (RM), parotid glands (PG), and lacrimal glands (LG). The activity of the RPT in each OAR was modeled to follow an exponential decay law governed by both the physical decay of the radionuclide (*λ*_phy_) and the organ-specific biological clearance rate (*λ*_r,X_):

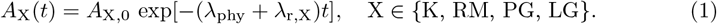

The same dynamics was assumed for tumor lesions (T) and for the remaining body (RB), defined as the part of the whole body (WB) that excludes OARs and tumor lesions, i.e., *M*_RB_ = *M*_WB_ −Σ_X_ *M*_X_, where X ∈ {K, RM, PG, LG, T}. The RB therefore represents a distinct compartment that receives any injected activity not taken up by the mCRPC tumor and OARs and clears it according to its own effective rate, *λ*_r,RB_.

The standardized uptake value (SUV) is defined as the image-derived activity concentration normalized by injected activity per body mass. Organ- and tumor-specific uptake fractions of the RPT were dynamically computed from their mean standardized uptake values (SUV_mean_) and corresponding masses (*M* ). Therefore, we have:

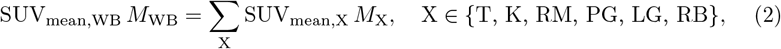

the mean standardized uptake value of the remaining body can be estimated from Eq.2 using its value in OARs and tumor as:

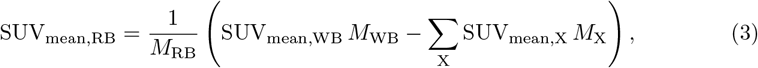

where X ∈ {K, RM, PG, LG, T}.

Finally, for each body compartment X (right panel in Fig 1), the activity fraction at the time of injection *t*_*i*_ was computed as

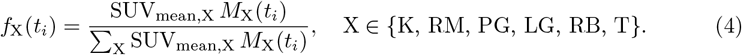

Note that Σ_X_ *f*_X_(*t*_*i*_) = 1. Also, *M*_T_(*t*_*i*_), the total tumor mass at the injection time *t*_*i*_, is the only time-dependent mass term on the right-hand side of Eq. 4.

In order to estimate overall survival for *in silico* patients, survival time was defined as the interval from treatment initiation until the total tumor mass reaches a critical threshold of 1000 g. Death was assumed to occur on average once this threshold was exceeded, based on clinical observations of tumor burden in advanced mCRPC [23].

### 2.2 Model equations

The system of ordinary differential equations (ODEs) describing tumor dynamics and radiopharmaceutical activity kinetics was formulated as follows:

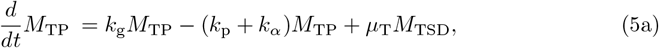

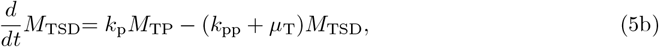

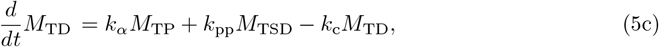

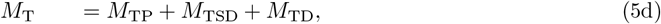

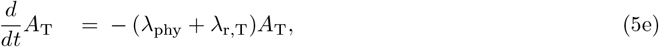

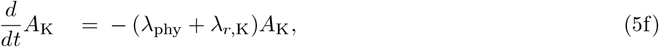

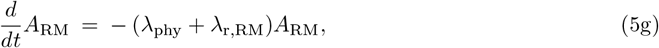

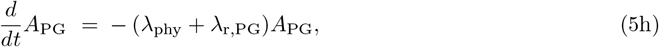

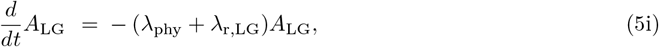

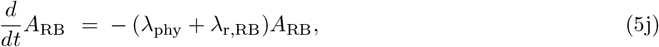

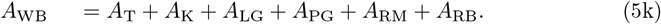

Eq. (5a) describes the dynamics of proliferating tumor cells (*M*_TP_). These cells grow at rate *k*_g_, transition to sublethally and lethally damaged states at activity-dependent rates *k*_p_ and *k*_*α*_, respectively, and receive repaired cells returning from the sublethal compartment at rate *µ*_T_. Eq. (5b) models the sublethally damaged tumor mass (*M*_TSD_), which is populated by radiation-induced damage from *M*_TP_ at rate *k*_p_, and either repairs back to *M*_TP_ at rate *µ*_T_ or progresses to lethal damage at rate *k*_pp_. The lethally damaged tumor mass (*M*_TD_), governed by Eq. (5c), accumulates cells transitioning from *M*_TP_ (at rate *k*_*α*_) and from *M*_TSD_ (at rate *k*_pp_), and is cleared from the system at rate *k*_c_. The total tumor mass (*M*_T_) is defined in Eq. (5d) as the sum of these three compartments.

The decay of radiopharmaceutical activity in the tumor (*A*_T_), kidneys (*A*_K_), red marrow (*A*_RM_), parotid glands (*A*_PG_), lacrimal glands (*A*_LG_), and remaining body (*A*_RB_) is described by Eqs. (5e)–(5j). In all cases, the activity is assumed to decline exponentially with an effective rate combining the physical decay constant *λ*_phy_ and the compartment-specific biological clearance rate *λ*_r,X_. The whole-body activity (*A*_WB_) is defined in Eq. (5k) as the sum of the activities of all body compartments.

### 2.3 Modeling activity injections

Activity administrations were modeled as instantaneous boluses, since the duration of RPT injection and initial biodistribution are negligible relative to the timescale of tumor and organ dynamics. The numerical integration of the ODE system (5) was stopped at each treatment time point *t*_*i*_ (*i* = 1, …, *n*, with *n* being the number of injections). At *t*_*i*_ the compartmental activities were updated to reflect the injected activity and its biodistribution, after which integration was resumed.

The injected activity is distributed among compartments according to their SUV_mean_-weighted mass fractions. For each OAR (X ∈ K, RM, PG, LG), as well as for the tumor and the remaining body, the fractional uptakes are calculated as in Eq. (4), evaluated with the current tumor mass *M*_T_(*t*_*i*_). The compartmental activities are then updated as

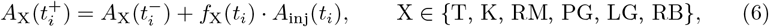

where *A*_inj_(*t*_*i*_) denotes the activity administered at time 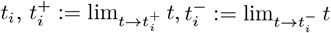, and *f*_X_(*t*_*i*_) are calculated as in Eq. (4). Eq. (6) ensures activity conservation across all compartments. No instantaneous changes are applied to the tumor mass compartments (*M*_TP_, *M*_TSD_, *M*_TD_) at the time of injection. Between consecutive administrations, all compartments evolve according to system (5).

### 2.4 Toxicity quantification

To quantify the radiobiological effect in OARs, we computed the biologically effective dose (BED), which accounts for total AD together with dose-rate and repair kinetics. Following Bodei *et al*. [24], the the BED for a specific OAR (BED_X_), was calculated as:

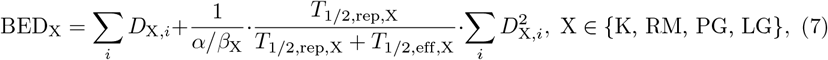

where *D*_X,*i*_ is the dose delivered to organ X during cycle *i, β*_X_*/α*_X_ is the ratio of X tissue-specific radiosensitivity parameters, *T*_1*/*2,rep,X_ is the sublethal-damage repair half-time in organ X, and *T*_1*/*2,eff,X_ is X organ-specific effective half-life of the radiopharmaceutical. Note that *T*_1*/*2,rep,X_ = log(2)*/µ*_X_ and *T*_1*/*2,eff,X_ = log(2)*/λ*_eff,X_.

At the individual patient level, toxicity was described using a simplified modeling approach: for each OAR X, a specific BED threshold 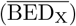 was set, representing the maximum cumulative BED above which RPT-induced toxicity is expected. We found in the literature BED threshold values for kidneys and lacrimal glands [25]. We estimated BED threshold values for red marrow and parotid glands using as *D*_*i*_ in Eq. (7) the value of AD from which severe toxicity is clinically observed under external beam radiation therapy, i.e., 2 Gy for red marrow [26] and 26 Gy for parotid glands [27]. Thus, if a VP accumulates a BED exceeding 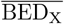 in a specific OAR X, toxicity is assumed to occur in that organ without distinguish between different severity of toxicity levels, although multiple thresholds could be implemented to predict graded toxic effects, as typically reported in clinical trials.

The population-level probability of toxicity (PoT) for a treatment regimen can be estimated using an *in silico* trial. The PoT is computed as the empirical fraction of VPs experiencing toxicity:

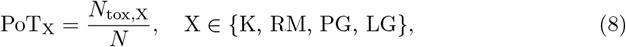

where *N*_tox,X_ is the number of VPs exceeding BED_X_ for organ X, and *N* is the total size of the virtual cohort. This approach provides a straightforward, sample-based estimate of the organ-specific probability of toxicity.

### 2.5 Model parameters

To capture the observed biological variability across patients [9], we first conducted an extensive review of the literature and experimental studies to identify biologically meaningful parameter ranges for our model. We extracted parameter ranges directly from published sources. The only parameter for which no information was available was *ε*, representing the probability of interaction between sublethal lesions. Therefore, the entire feasible interval (0, 1) was explored to cover all scenarios.

The parameters are organized into three categories, each summarized in a separate table. Table 1 presents stochastic parameters for which literature-based ranges are available. Table 2 compiles constant parameters, for which literature-based ranges are not available to the best of our knowledge. Finally, Table 3 contains parameters that were derived from analytical *formulae*. For detailed parameter explanations, see the Supplementary Section S1.

**Table 1.** Stochastic parameters used in the model, including their distributions, ranges, and units. Parameters include the Biological Clearance Rate (BCR) and the coefficients of the Linear-Quadratic (LQ) model.

| Symbol | Description | Range | Unit | Ref. |
| --- | --- | --- | --- | --- |
| $M_T(0)$ | Initial tumor mass | (0, 100) | g | [28] |
| $M_K$ | Kidney mass | (231, 503) | g | [29] |
| $M_{PG}$ | Parotid gland mass | (30, 60) | g | [30] |
| $M_{LG}$ | Lacrimal gland mass | (1, 1.56) | g | [31] |
| $M_{RM}$ | Mass of red marrow | (1413, 2937) | g | [32] |
| $\alpha$ | Linear coefficient in the LQ model | (0.01, 0.15) | Gy<sup>-1</sup> | [33] |
| $\beta$ | Quadratic coefficient in the LQ model | (0.003, 0.058) | Gy<sup>-2</sup> | [33] |
| $k_c$ | Tumor clearance constant | (0.0001, 0.001) | h<sup>-1</sup> | [34] |
| $k_g$ | Tumor growth constant for mCRPC | (0.0001, 0.0019) | h<sup>-1</sup> | [35] |
| $\lambda_{r,T}$ | BCR for the tumor | (10<sup>-5</sup>, 0.045) | h<sup>-1</sup> | [36] |
| $\lambda_{r,K}$ | BCR for the kidneys | (0.004, 0.032) | h<sup>-1</sup> | [36] |
| $\lambda_{r,PG}$ | BCR for parotid glands | (0.0117, 0.0303) | h<sup>-1</sup> | [36] |
| $\lambda_{r,RM}$ | BCR for red marrow | (0.006, 0.077) | h<sup>-1</sup> | [37] |
| $\lambda_{r,RB}$ | BCR for the remaining body | (0.0032, 0.0286) | h<sup>-1</sup> | [36] |
| $\alpha/\beta_K$ | Radiobiological $\alpha/\beta$ for kidneys | (1.4, 4.3) | Gy | [38] |
| $\varepsilon$ | Sublethal lesions interaction probability | (0, 1) | – | – |
| $\mu_K$ | Sublethal damage repair rate in kidneys | (0.1, 0.57) | h<sup>-1</sup> | [38] |
| $M_{WB}$ | Mass of the whole body | (6 · 10<sup>4</sup>, 10<sup>5</sup>) | g | [39] |
| $SUV_{mean,T}$ | Mean SUV in tumor | (3, 80) | – | [23] |
| $SUV_{mean,PG}$ | Mean SUV in parotid gland | (5, 16) | – | [40] |
| $SUV_{mean,LG}$ | Mean SUV in lacrimal gland | (4.8, 8.4) | – | [40] |
| $SUV_{mean,K}$ | Mean SUV in kidneys | (4, 30) | – | [40] |
| $SUV_{mean,RM}$ | Mean SUV in red marrow | (0.3, 1.3) | – | [40] |

**Table 2.** Constant parameters used in the model, together with their fixed values, units, and corresponding references.

| Symbol | Description | Value | Unit | Ref. |
| --- | --- | --- | --- | --- |
| $E_{e,^{177}\text{Lu}}$ | Energy emitted per $\beta$ decay ($^{177}\text{Lu}$) | 0.0849 | mJ·MBq<sup>-1</sup>·h<sup>-1</sup> | [41] |
| $S_{\gamma,WB}$ | Self-dose S-value for whole-body $\gamma$ rays | 1.85 · 10<sup>-6</sup> | Gy·Kg<sup>2/3</sup>·MBq<sup>-1</sup>·h<sup>-1</sup> | [42] |
| $SUV_{mean,WB}$ | Mean SUV in whole body | 1 | – | [43] |
| $\lambda_{phy}$ | Physical decay constant of $^{177}\text{Lu}$ | 0.0044 | h<sup>-1</sup> | [41] |
| $\mu_T$ | Sublethal damage repair rate in tumor | 0.365 | h<sup>-1</sup> | [44] |
| $\alpha/\beta_{PG}$ | Radiobiological $\alpha/\beta$ for PG | 4.4915 | Gy | [45] |
| $\mu_{PG}$ | Sublethal damage repair rate in PG | 0.462 | h<sup>-1</sup> | [45] |
| $\alpha/\beta_{RM}$ | Radiobiological $\alpha/\beta$ for RM | 15 | Gy | [45] |
| $\mu_{RM}$ | Sublethal damage repair rate in RM | 0.4621 | h<sup>-1</sup> | [45] |
| $T_{1/2,rep,LG}$ | Repair half-time for LG | 1.5 | h | [25] |
| $\lambda_{r,LG}$ | BCR for lacrimal glands | 0.0204 | h<sup>-1</sup> | [46] |
| $\alpha/\beta_{LG}$ | Radiobiological $\alpha/\beta$ for lacrimal glands | 0.222 | Gy | [25] |
| $\overline{BED}_K$ | BED threshold for toxicity in K | 39 | Gy | [25] |
| $\overline{BED}_{LG}$ | BED threshold for toxicity in LG | 68 | Gy | [25] |
| $\overline{BED}_{RM}$ | BED threshold for toxicity in RM | 2.02 | Gy | [24] |
| $\overline{BED}_{PG}$ | BED threshold for toxicity in PG | 31.5 | Gy | [24] |

**Table 3.** Summary of model parameters expressed through analytical *formulae*, including their biological meaning, mathematical expression, units, and corresponding references.

| Symbol | Description | Formula | Unit | Ref. |
| --- | --- | --- | --- | --- |
| $\lambda_{\text{eff},X}$ | Effective release from tissue X | $\lambda_{r,X} + \lambda_{\text{phy}}$ | $\text{h}^{-1}$ | [17] |
| $A_{\text{cum},X}$ | Cumulated activity in tissue X | $\int A_X dt$ | $\text{MBq}\cdot\text{h}$ | [17] |
| $D_X$ | Dose rate to organ X | $S_X \times A_{\text{cum},X}$ | Gy | [47] |
| $S_X$ | S-value of organ X | $E_{e,^{177}\text{Lu}}/M_X$ | $\text{Gy}\cdot\text{MBq}^{-1}\cdot\text{h}^{-1}$ | [47] |
| $\dot{D}_T$ | Dose rate to tumor | $S_T \times A_T$ | $\text{Gy}\cdot\text{h}^{-1}$ | [47] |
| $S_T$ | S-value of tumor | $E_{e,^{177}\text{Lu}} \times (M_T)^{-0.99}$ | $\text{Gy}\cdot\text{MBq}^{-1}\cdot\text{h}^{-1}$ | [17] |
| $k_\alpha$ | Direct lethal damage rate | $\alpha \times \dot{D}_T$ | $\text{h}^{-1}$ | [17] |
| $k_p$ | Sub-lethal damage rate | $2p \dot{D}_T$ | $\text{h}^{-1}$ | [17] |
| $k_{pp}$ | Interaction rate of sub-lethal and lethal damage | $(p + \alpha) \varepsilon \dot{D}_T$ | $\text{h}^{-1}$ | [17] |
| $p$ | Yield of sublethal lesions per unit AD | $\sqrt{\beta/\varepsilon}$ | $\text{Gy}^{-1}$ | [17] |
| $D_{\text{RM}}$ | AD to red marrow | $S_{\text{RM}} A_{\text{cum},\text{RM}} + S_{\gamma,\text{WB}}/(M_{\text{WB}})^{2/3} A_{\text{cum},\text{WB}}$ | Gy | [41] |

### 2.6 Sobol Indices

To quantify how uncertainty in the model parameters affects the outputs of the nonlinear system (5), we constructed a polynomial chaos expansion surrogate and computed first-order and total-order Sobol indices. Only the parameters from Table 1 that appear explicitly in the equations were included, ensuring that no artificial variability was introduced from quantities not influencing the system dynamics. All uncertain parameters were modeled as independent uniform distributions with their biologically plausible ranges reported in Table 1.

The surrogate was trained using 10^4^ Latin hypercube samples. Let ***θ*** = (*θ*_1_, *θ*_2_, …, *θ*_*d*_) denote the vector of uncertain model parameters considered in the sensitivity analysis. Because the model contains nonlinear proliferation–decay couplings, multiplicative interactions between tumor compartments, and sequential elimination processes across multiple organs, the corresponding solution map

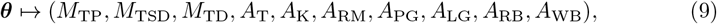

exhibits strong curvature and higher-order parameter interactions. Polynomial chaos theory shows that when the underlying dynamics include such nonlinearities and cross-interactions, low-order expansions tend to under-represent the geometry of the response surface, leading to biased or unstable sensitivity estimates. High-degree polynomial bases are therefore recommended in nonlinear ODE systems with coupled components [48, 49, 50, 51]. For this reason, we adopted a sixth-order polynomial basis, which provides sufficient expressive power to approximate the dependence of all model outputs on ***θ***.

To avoid the combinatorial explosion typically associated with high-order expansions, we employed sparse regression via least angle regression, as implemented in UQLab [52, 48]. This approach automatically selects only the significant polynomial modes, ensuring that the high-order polynomial chaos expansion remains computationally tractable while retaining accuracy. The clinical treatment schedule–six administrations of 7.4 GBq of [^177^Lu]Lu-PSMA-617 every six weeks–was encoded directly in the ODE system and consistently represented in the surrogate.

### 2.7 Virtual patients and *in silico* trials

A key strength of the proposed model is its ability to generate VPs, thereby enabling *in silico* trials to be conducted. A VP is a computational construct representing an individual subject through the combination of biologically and physiologically plausible parameter values. Unlike generic model instances, each VP captures a distinct set of biological and pharmacokinetic features that together govern tumor growth, radiopharmaceutical kinetics and organ-specific ADs.

VPs were generated based on the parameter ranges summarized in Table 1. These ranges reflect the variability observed between patients in biological data, ensuring that not all patients share the same characteristics. As the exact statistical distributions of many parameters are unknown, uniform distributions were adopted across the reported physiological intervals, in line with prior modeling studies [53, 54]. Each VP was obtained by randomly selecting a value for each parameter within the prescribed ranges.

By repeating this process *N* times, we obtain a virtual cohort of *N* patients. Simulating their individual responses under a given treatment protocol allowed us to perform *in silico* trials. As far as the model reflects the right biology and parameter estimates are correct, these trials enabled the evaluation of population-level outcomes, inter-patient variability, and optimization of dosing strategies, all achieved without posing any risk to actual patients or compromising the success of real clinical trials [12].

Moreover, calibrating model parameters with patient-specific data could enable the generation of a digital twin and the exploration of personalized treatments [55, 56, 57].

### 2.8 Simulation details

Before the first administration, neither RPT activity nor RPT-induced damage were present; consequently, the initial tumor mass was assumed to consist solely of proliferating cells, *M*_T_(0) = *M*_TP_(0), with *M*_TSD_(0) = *M*_TD_(0) = 0, and *A*_X_(0) = 0, X ∈ {K, RM, PG, LG, T, RB} [17]. Thus, if the first cycle was applied at *t*_*i*_ (possibly *t*_*i*_ = 0), the initial conditions for the activities satisfied 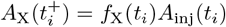 for each compartment X ∈ {K, RM, PG, LG, T, RB}, corresponding to Eq. (6) with *t*_*i*_ and 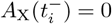. Subsequently, the kinetics evolved according to Eqs. (5e)–(5j).

Simulations were run until they reached predefined endpoints, which were determined by the biological context or the experimental design. These endpoints may correspond to specific time points or a tumor reaching a critical size threshold, such as complete remission, recurrence, or lethal burden.

The model equations were solved using the variable step ODE solver `ode45` included in the scientific software package `MATLAB` (R2023b, The MathWorks, Inc., Natick, MA, USA), running on a Mac Studio with an Apple M1 Ultra chip and 128 GB of RAM.

## 3 Results

### 3.1 Model validation

In order to evaluate the reliability and predictive capacity of our mathematical framework, we conducted a validation process in which model outputs were compared with clinical studies reported in the literature. Specifically, we implemented *in silico* treatment schemes simulating those described in the clinical studies by Violet *et al*. [18] and Scarpa *et al*. [19], applying the same treatment protocols as in the original trials to comparable VP cohorts.

In the clinical trial reported by Violet *et al*. [18], 30 patients received an average of four treatment cycles, each consisting of 7.8 GBq, administered at six-week intervals. This treatment scheme was implemented computationally to reproduce the clinical study as closely as possible. The mean AD ± standard deviation reported per administered activity (Gy/GBq) was 0.39 ± 0.15 for kidneys, 0.58 ± 0.43 for parotid glands, 0.36 ± 0.18 for lacrimal glands, 0.11 ± 0.10 for red marrow, and 2.3 ± 1.5 for tumor lesions [18]. In comparison, our *in silico* cohort yielded corresponding mean AD values of 0.69 *±* 0.43 for kidneys, 0.43 *±* 0.19 for parotid glands, 0.27 *±* 0.07 for lacrimal glands, 0.036 *±* 0.0194 for red marrow, and 1.75 *±* 1.15 for tumor lesions.

Scarpa *et al*. [19] evaluated 10 patients who received two or three treatment cycles, with an administered activity of 6.1 ± 0.3 GBq per cycle. The mean AD ± standard deviation reported per administered activity (Gy/GBq) was 0.60 ± 0.362 for kidneys, 0.042 ± 0.028 for red marrow, 0.561 ± 0.248 for parotid glands, and 1.006 ± 0.69 for lacrimal glands. For tumor lesions, the mean AD averaged across all patients was found to be 2.8 ± 0.5 Gy/GBq [19]. By contrast, our *in silico* cohort yielded corresponding mean AD values of 1.22 ± 0.66 for kidneys, 0.32 ± 0.05 for parotid glands, 0.29 ± 0.08 for lacrimal glands, 0.03 ± 0.013 for red marrow, and 3.37 ± 2.27 for tumor lesions.

In Fig 2(a), the mean ADs and their standard deviations are shown for the various OARs, both from the clinical study [18] and from the *in silico* trial performed using our mathematical model. Although all confidence intervals overlap, the closest agreement is observed in the mean AD to the tumor mass. Similarly, Fig 2(b) presents the mean ADs and corresponding standard deviations for the clinical data [19] and for our simulations across the different OARs. In this case, the mean ADs for the red marrow and tumor show good agreement, whereas a larger deviation is observed for the lacrimal glands.

**Fig 2.**
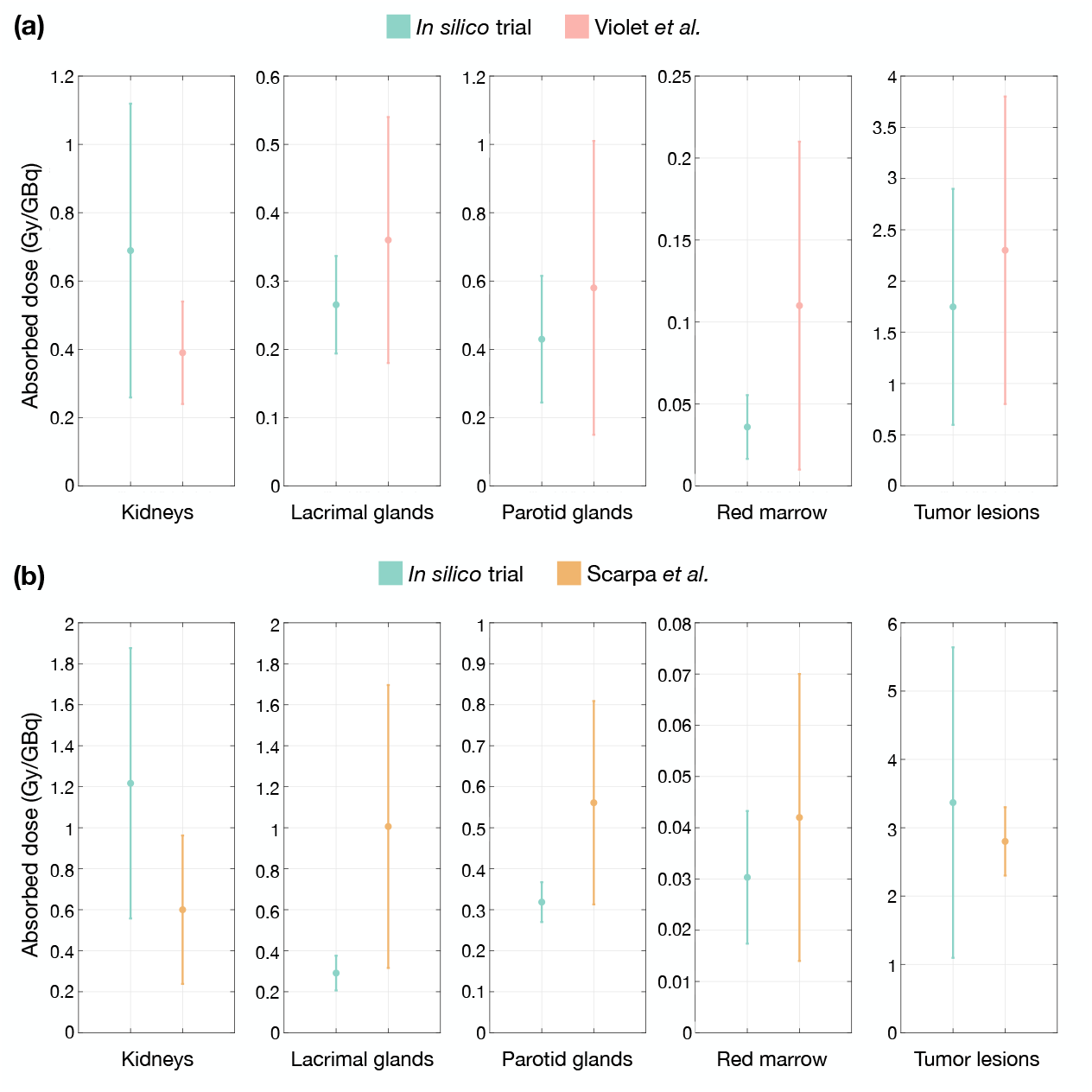
Comparison of ADs (Gy/GBq) in the main OARs (kidneys, red marrow, parotid glands, and lacrimal glands) and tumor obtained from the mathematical model using two clinical trials. Panel (a) shows the comparison with the study by Violet *et al*. [18], while Panel (b) shows the comparison with the study by Scarpa *et al*. [19].

In addition to validating the AD in different OARs, we aimed to assess our model’s ability to predict patient survival. To validate these results, we considered the TheraP clinical trial [20, 58, 59], which compared [^177^Lu]Lu-PSMA-617 and cabazitaxel in men with mCRPC that had progressed after docetaxel treatment. We focused exclusively on the cohort treated with [^177^Lu]Lu-PSMA-617. These patients received up to six treatment cycles, administered every six weeks, starting at 8.5 GBq and decreasing by 0.5 GBq per cycle down to a minimum of 6.0 GBq. The trial reported a median follow-up of 35.7 months, during which 200 patients were randomized, with 99 assigned to [^177^Lu]Lu-PSMA-617 and 101 to cabazitaxel [20].

Based on these data, we put forward an *in silico* trial to match the same number of VPs as the clinical trial [^177^Lu]Lu-PSMA-617 arm, involving 99 patients. This design allowed us for a direct comparison of overall survival outcomes.

In Fig 3 (a) we present the restricted mean survival time up to 35.7 months (the follow-up period used in the TheraP study [20]), for both the clinical and *in silico* trials. In TheraP [20], the reported restricted mean survival time for the [^177^Lu]Lu-PSMA-617 arm was 19.6 months (95% confidence interval (CI): 17.4–21.8). Our simulation yielded a restricted mean survival time of 19.47 months (95% CI: 15.55–23.40). Fig 3 (b) shows the corresponding Kaplan–Meier survival curves. A log-rank test detected no statistically significant difference between survival distributions (two-sided *p* = 0.63), indicating that, within follow-up, the *in silico* and clinical outcomes were statistically indistinguishable.

**Fig 3.**
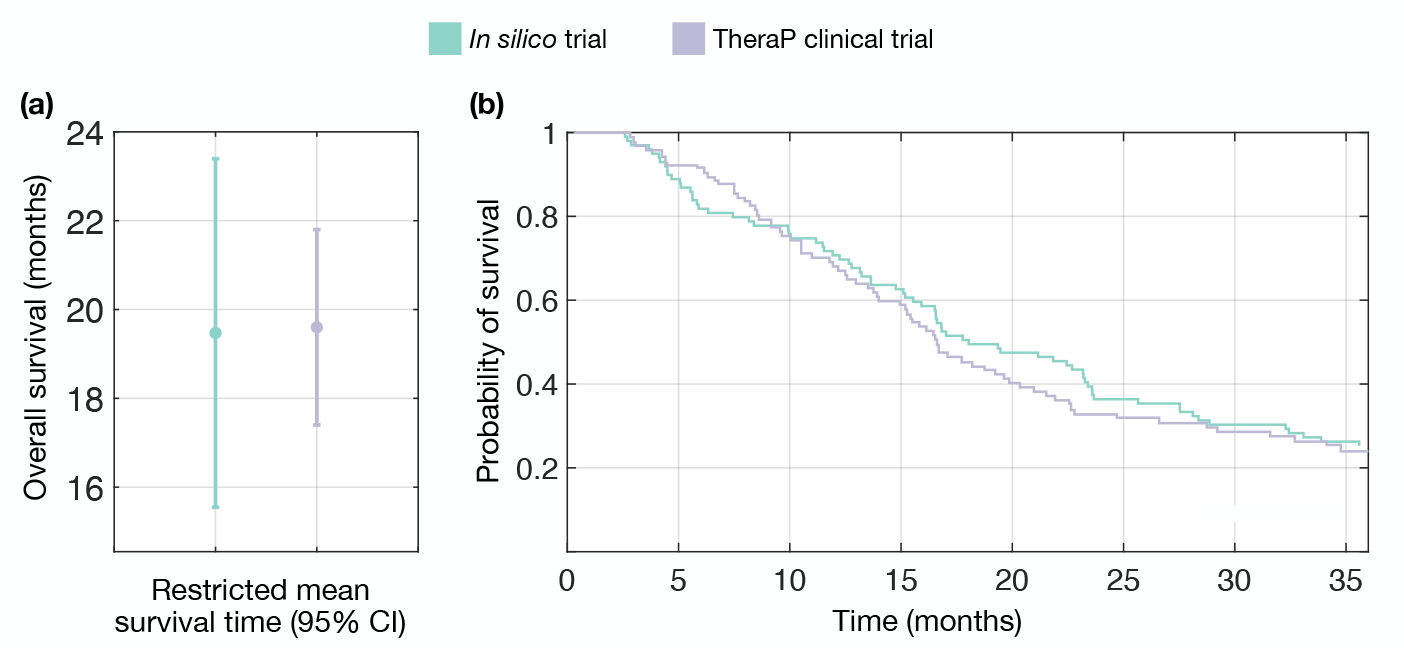
**(a)** Restricted mean survival time up to 35.7 months, obtained from the *in silico* model and compared with results from the TheraP clinical trial [20]. The markers indicate the estimated restricted mean survival time, and the error bars represent the corresponding 95% confidence intervals (CIs). **(b)** Kaplan–Meier survival curves comparing the *in silico* model with the TheraP clinical trial (log-rank *p* = 0.63) [20].

For additional validation, we considered the LuPSMA clinical trial [21]. In this study, 30 patients with [^177^Lu]Lu-PSMA-617-eligible mCRPC who had received at least one prior line of chemotherapy were enrolled. The mean administered radioactivity was 7.5 GBq per cycle. In our *in silico* trial, we assumed that each VP received a total of four cycles (in the LuPSMA trial some patients received fewer cycles). The reported median follow-up was 25 months.

Figure 4 (a) displays the median overall survival (mOS) and the corresponding 95% CI for both the clinical and *in silico* trials. Unlike in the TheraP validation [20], where the restricted mean survival time was reported, here the LuPSMA study [21] provides median survival as the primary summary statistic. In this clinical trial, the mOS was 13.5 months (95% CI: 10.4–22.7). Our *in silico* trial yielded a closely comparable estimate of 14.62 months (95% CI: 7.50–23.25).

**Fig 4.**
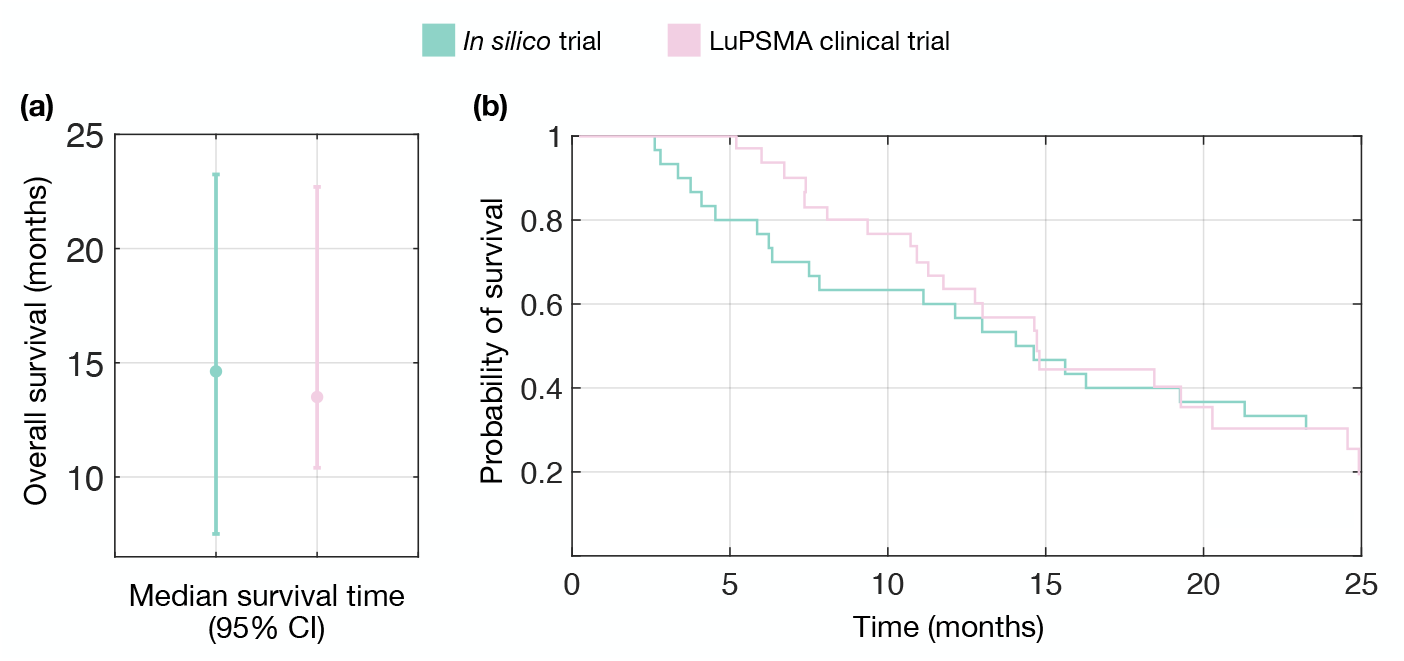
**(a)** Median survival time at 25 months obtained from the *in silico* model and compared with results from the LuPSMA clinical trial [21]. The markers indicate the estimated restricted mean survival time, and the error bars represent the corresponding 95% confidence intervals (CI). **(b)** Kaplan–Meier survival curves comparing the *in silico* model with the LuPSMA clinical trial (log-rank *p* = 0.29) [21].

Figure 4 (b) shows the Kaplan–Meier survival curves derived from the *in silico* trial and the LuPSMA study [21]. As in the previous case, a log-rank test indicated no significant difference between survival distributions (*p* = 0.29), supporting that the survival outcomes predicted by our *in silico* model are consistent with those observed in LuPSMA [21].

Note that the supplementary material includes an Excel file named *ParametersVP*.*xlsx*, which contains tables detailing the parameter values for each VP cohort generated to validate the model. These tables list the set of parameters drawn for each VP and used in the simulation of our *in silico* trial.

### 3.2 Assessing virtual cohort size for population-level reliability

A critical question when generating a VP cohort, as in recruiting real patient cohorts, is whether the number of VPs chosen is large enough to provide faithful results matching those of the *N*→ ∞ limit. To address this question, we performed an empirical convergence analysis based on Kaplan-Meier survival estimates, inspired by the strategy presented in [60]. For increasing cohort sizes *N* we generated 10 independent virtual cohorts sampled from the same parameter ranges (Table 1). We then conducted pairwise comparisons between the survival curves of each cohort under standard treatment (six 6-week cycles of 7.4 GBq each one) using two-sided log-rank tests, yielding 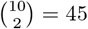 pairwise tests for each *N* (= 50, 100, … ). The smallest cohort size *N* at which no pairwise comparisons showed statistically significant differences at the 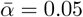 level was *N* = 500 (for *N <* 500, at least one comparison was significant; see Fig 5, where p-values are sorted in ascending order to facilitate visualization and comparison between rows). Note that with 10 cohorts (*m* = 45 tests per *N*), the expected number of false positives (Type I errors) under the null hypothesis is 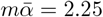, and the probability of observing at least one false positive is 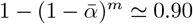. Thus, isolated rejections observed for smaller *N* are compatible with chance findings due to multiplicity rather than systematic differences. Requiring zero rejections is therefore a conservative convergence criterion, and it is first satisfied at a cohort size of *N* = 500.

**Fig 5.**
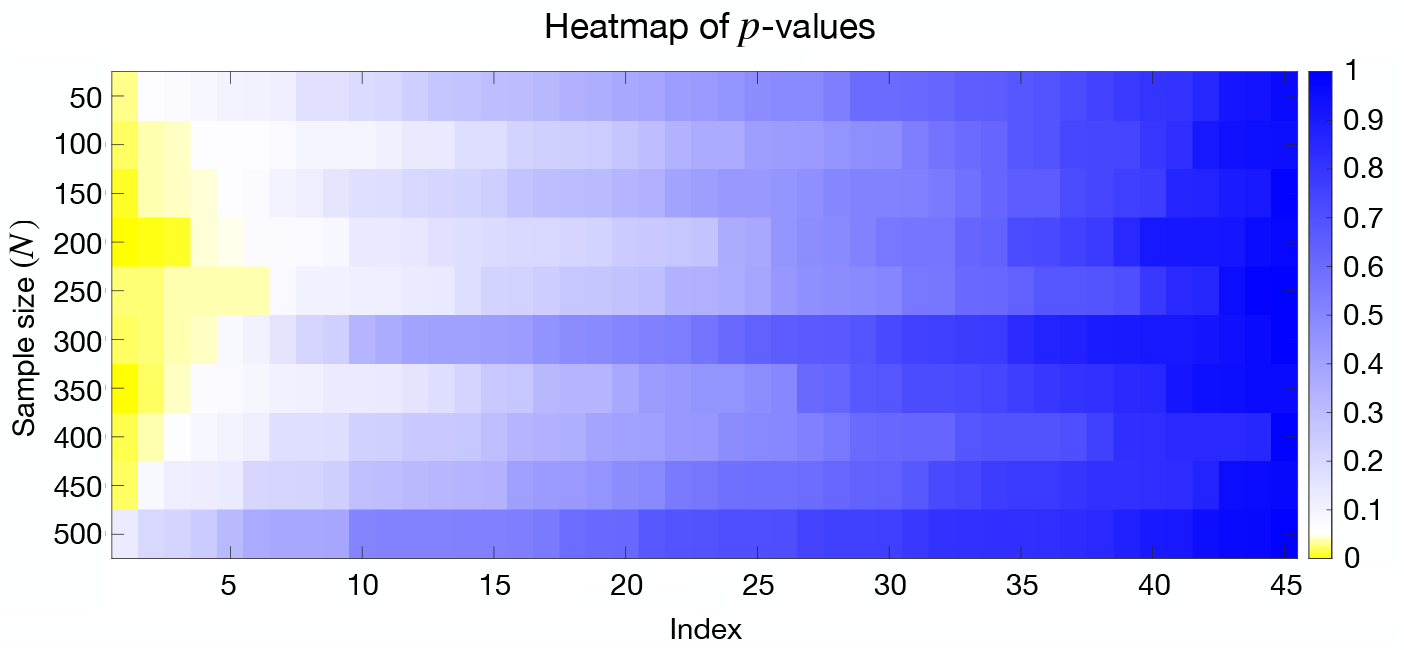
Heatmap of *p*-values sorted in ascending order for different virtual patient cohort sizes (*N*). Each row corresponds to a cohort size (from *N* = 50 to *N* = 500), and each column represents a pairwise comparison index. Colors indicate the magnitude of *p*-values, with yellow for statistically significant difference (*p*-values *<* 0.05), blue for larger *p*-values, and white centered at *p*-value= 0.05. This visualization highlights how statistical significance varies with cohort size, supporting the choice of *N* = 500 for stable and reproducible results.

These results indicate that a cohort size of *N* = 500 VPs provides stable and reproducible population-level outcomes: inter-cohort variability in Kaplan-Meier curves is no larger than the one expected by chance, supporting the generalizability of conclusions drawn from a single simulated cohort of *N* = 500 while retaining computational efficiency.

### 3.3 Investigating cycle dosages

Once the model was validated and the virtual cohort size for population-level reliability found, we used the model to investigate the effect of injection fractionation on standard RPT. To do so, we performed *in silico* trials using a virtual cohorts of 500 VPs to ensure population-level reliability, as demonstrated in Section 3.2. Specifically, we simulated a series of treatment regimens by varying the number of injections (*n*_inj_) while keeping the total administered activity fixed at 44.4 GBq and the cycle length at 6 weeks, consistent with clinical practice. Fig 6 and Supplementary Table S1 summarizes the results, including the mOS and the probability of toxicity (PoT) for OARs. Unless otherwise specified, doses were split equally across injections (44.4 GBq/*n*_inj_). regimens marked with an asterisk (*) were simulated with asymmetric dose distributions with the specific split being reported in Supplementary Table S1.

**Fig 6.**
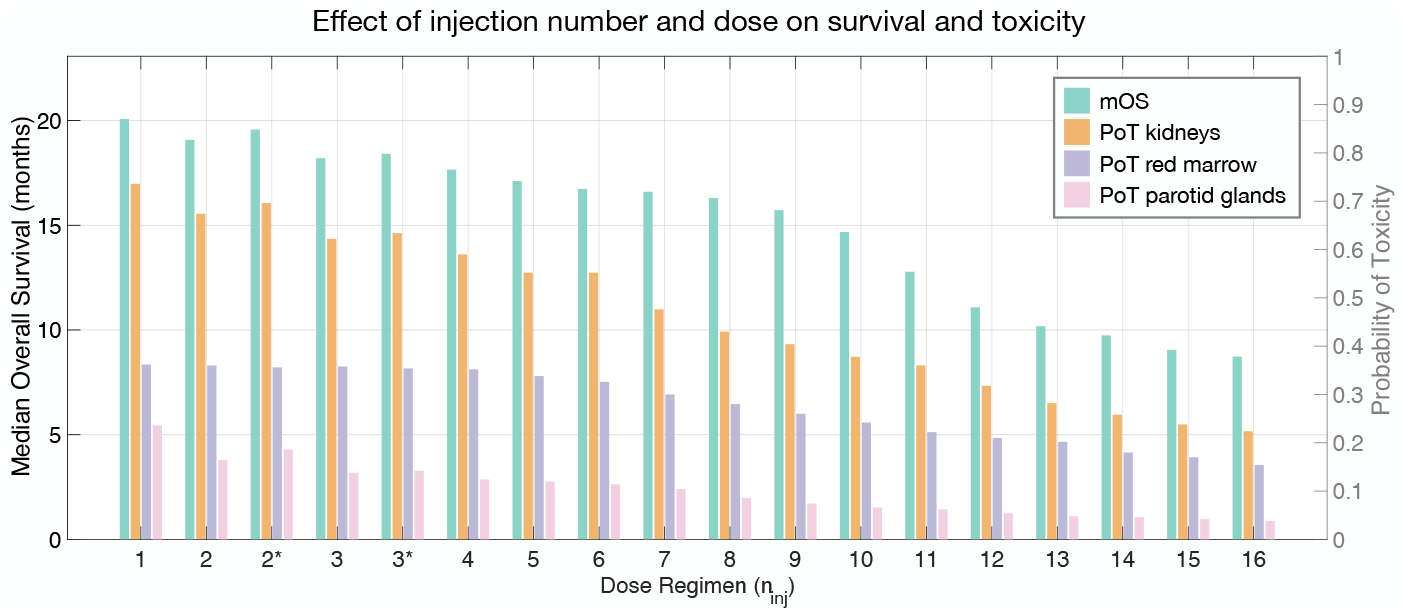
Summary of median overall survival (mOS) outcomes (left y-axis) and probabilities of toxicity (PoT) for organs-at-risk (right y-axis) under regimens with varying number of injections. Each regimen assumes a fixed total activity (44.4 GBq) and a 6-week administration cycle length; regimens marked with * use a heterogeneous (asymmetric) dose distribution.

The mOS improved with regimens featuring fewer, higher-activity injections. The highest mOS was obtained with the one-injection schedule (20.07 months), followed by the asymmetric (19.57 months) and symmetric (19.08 months) two-injection protocols, corresponding to gains of approximately 20%, 17%, and 14%, respectively, relative to the standard protocol (16.73 months). However, none of these regimens achieved a statistically significant improvement (the smallest *p*-value was 0.056 for the one-injection regimen). Regimens with up to eight injections maintained a mOS larger than 16 months. However, increasing the number of injections to 12 or more led to a marked decrease in mOS–around 11 months with 12 injections and down to 8.73 months with 16 injections–despite identical total administered activity.

Survival improvements observed in the low-to-moderate fractionation range were accompanied by increased renal and parotid toxicity: the fewer the injections, the higher the probability of toxicity. The one-injection protocol yielded the highest PoT_K_ (0.736) and PoT_PG_ (0.236), both showed a decreasing trend with increasing *n*_inj_, reaching 0.224 and 0.038 at 16 injections, respectively. Red marrow toxicity exhibited less sensitivity to injection number, with PoT_RM_ remaining relatively stable up to seven injections (0.3–0.362) and gradually declining thereafter to 0.154 at 16 injections. Lacrimal gland toxicity was negligible in all cases, being observed only in the one-injection schedule (0.02).

Asymmetric dosing schedules emerged as a promising strategy. The 2* schedule (7.4 and 37.0 GBq) resulted among the best survival outcomes (mOS 19.57 months) while incurring only modest increases in PoT_K_ and PoT_PG_ relative to its symmetric two-injection counterpart. Similarly, the 3* schedule slightly outperformed its symmetric regimen in mOS (18.41 vs. 18.20 months) with minimal additional toxicity.

Among the tested regimens, the 2* asymmetric schedule offered a favorable efficacy-toxicity trade-off. While the one-injection protocol maximized survival, elevated renal and parotid glands toxicities (and, rarely, lacrimal toxicity) may limit its feasibility. For more conservative scenarios prioritizing toxicity reduction, the eight-injection schedule provided a reasonable compromise (mOS of 16.30 months; PoT_K_ of 0.43; PoT_RM_ of 0.28; PoT_LG_ of 0; PoT_PG_ of 0.086).

### 3.4 Investigating cycle lengths

To explore the effect of treatment timing on the standard RPT schedule, we performed *in silico* trials using the same virtual cohort of 500 VPs as in Section 3.3. This approach ensures robust population-level results (see Section 3.2) and direct comparison with the results presented in Section 3.3. Specifically, we simulated a series of regimens with varying cycle lengths, ranging from 0.125 (approximately one day) to 16 weeks, while keeping both the number of injections (*n*_inj_ = 6) and the dose per-injection (7.4 GBq) constant. This setup leads to the same total administered activity of 44.4 GBq across all schedules. Fig 7 and Supplementary Table S2 collect the resulting mOS and predicted PoT for the considered OARs.

**Fig 7.**
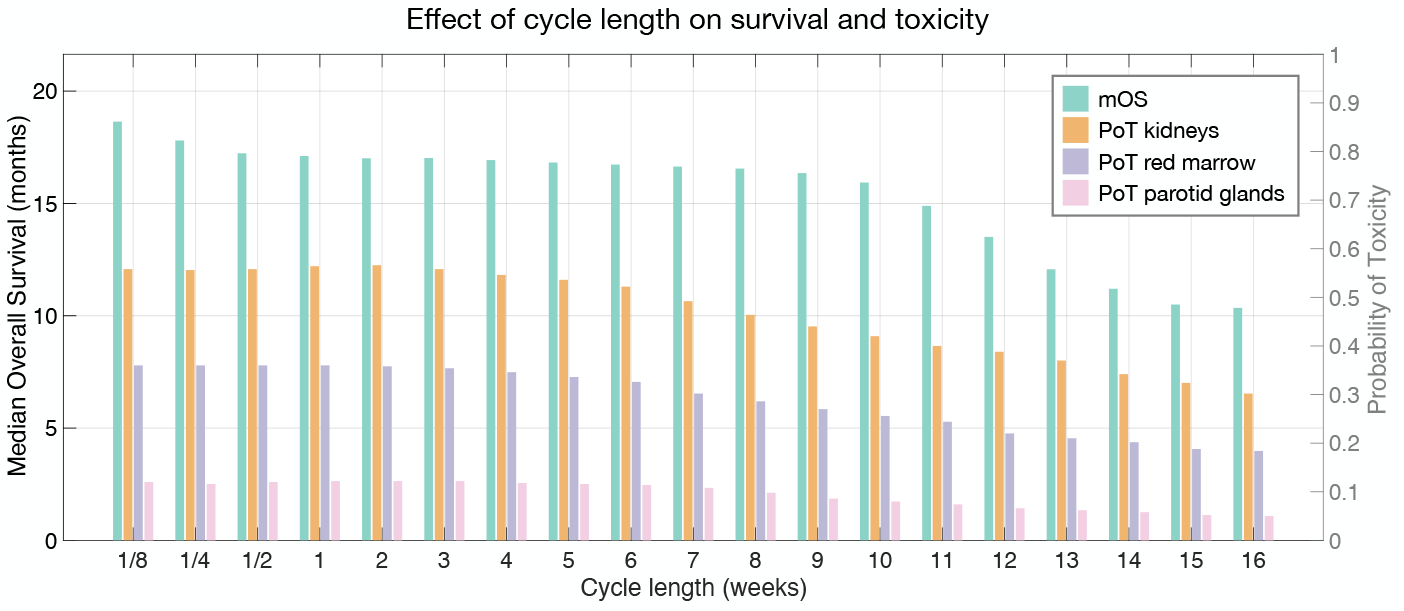
Summary of median overall survival (mOS) outcomes (left y-axis) and probabilities of toxicity (PoT) for organs-at-risk (right y-axis) under regimens varying the cycle length. Each regimen assumes six injections of 7.4 GBq per injection (total 44.4 GBq).

The mOS improved with regimens featuring shorter cycle length. The highest mOS (18.64 months) occurred at the shortest interval (0.125 weeks), with only a modest gain (≈11%) relative to the standard 6 week cycle (mOS = 16.73 months). Across a broad range of cycle lengths, particularly between 2 and 9 weeks, mOS remained relatively stable, hovering around 16.75 months. Very short cycles (*<* 2 weeks) also yielded similar outcomes (mOS *>* 17 months). By contrast, extending the cycle length to ≥ 12 weeks led to a sharp decline in survival: at ≥12 weeks, mOS dropped to 13.51 months and decreased further to 11.20 and 10.35 months for 14 and 16 weeks, respectively.

Toxicity varied modestly across the evaluated schedules. Renal, parotid, and red marrow probabilities of toxicity remained within a relatively narrow range up to six weeks (PoT_K_ ∈ [0.52, 0.56], PoT_PG_ ∈ [0.11, 0.12], PoT_RM_ ∈ [0.32, 0.36]). Beyond this point, toxicity progressively decreased with increasing cycle length, reaching PoT_K_ = 0.302, PoT_PG_ = 0.05 and PoT_RM_ = 0.184 at sixteen weeks. No toxicity was observed in the lacrimal glands for any of the cycle lengths evaluated (*n*_inj_ = 6, 7.4 GBq per injection).

We further compared the standard 6 week cycle with the 9 week cycle because they achieved similar mOS, while the latter regimen resulted in lower probabilities of toxicity. Figure 8 shows boxplots of survival and OAR BEDs for the cohort of 500 VPs under the two regimens. We found no significant differences in survival, whereas the BEDs in all OARs were statistically different and lower under the 9 week regimen.

**Fig 8.**
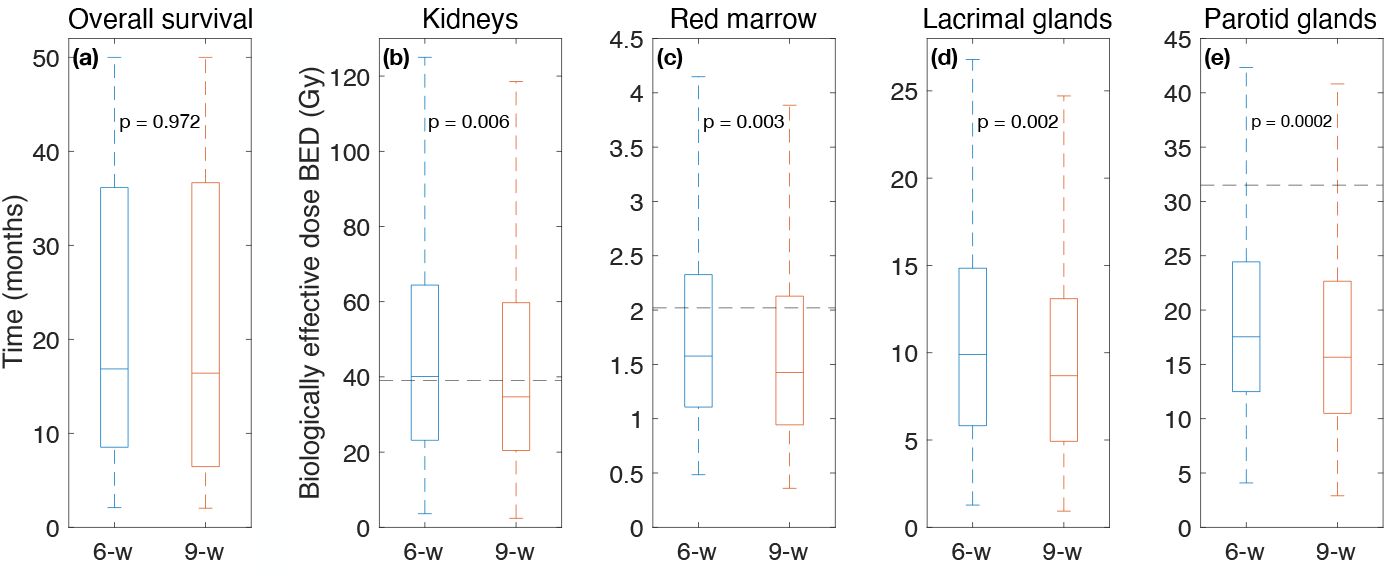
Comparison of (a) overall survival, (b) kidney toxicity, (c) red marrow toxicity, (d) lacrimal glands toxicity, and (e) parotid glands toxicity, between the standard (6-w) and 9 week (9-w) regimens 500 virtual patients were simulated in each branch of the comparison with dose per-injection of 7.4 GBq. P indicates the p-values for log-rank test (overall survival) and rank-sum tests (BEDs in organs-at-risk). Horizontal dotted black lines represent organ BED thresholds.

### 3.5 Sensitivity analysis

Fig 9 shows the first-order and total-order Sobol indices for tumor populations and activity. The remaining figures for the other variables are provided in Supplementary Figures. In general, the parameters with the most substantial influence according to first-order Sobol indices (*S*_1_) matched those most influential in total-order indices (*S*_*T*_ ), although the *S*_1_ values were systematically lower than *S*_*T*_ [61, 62].

**Fig 9.**
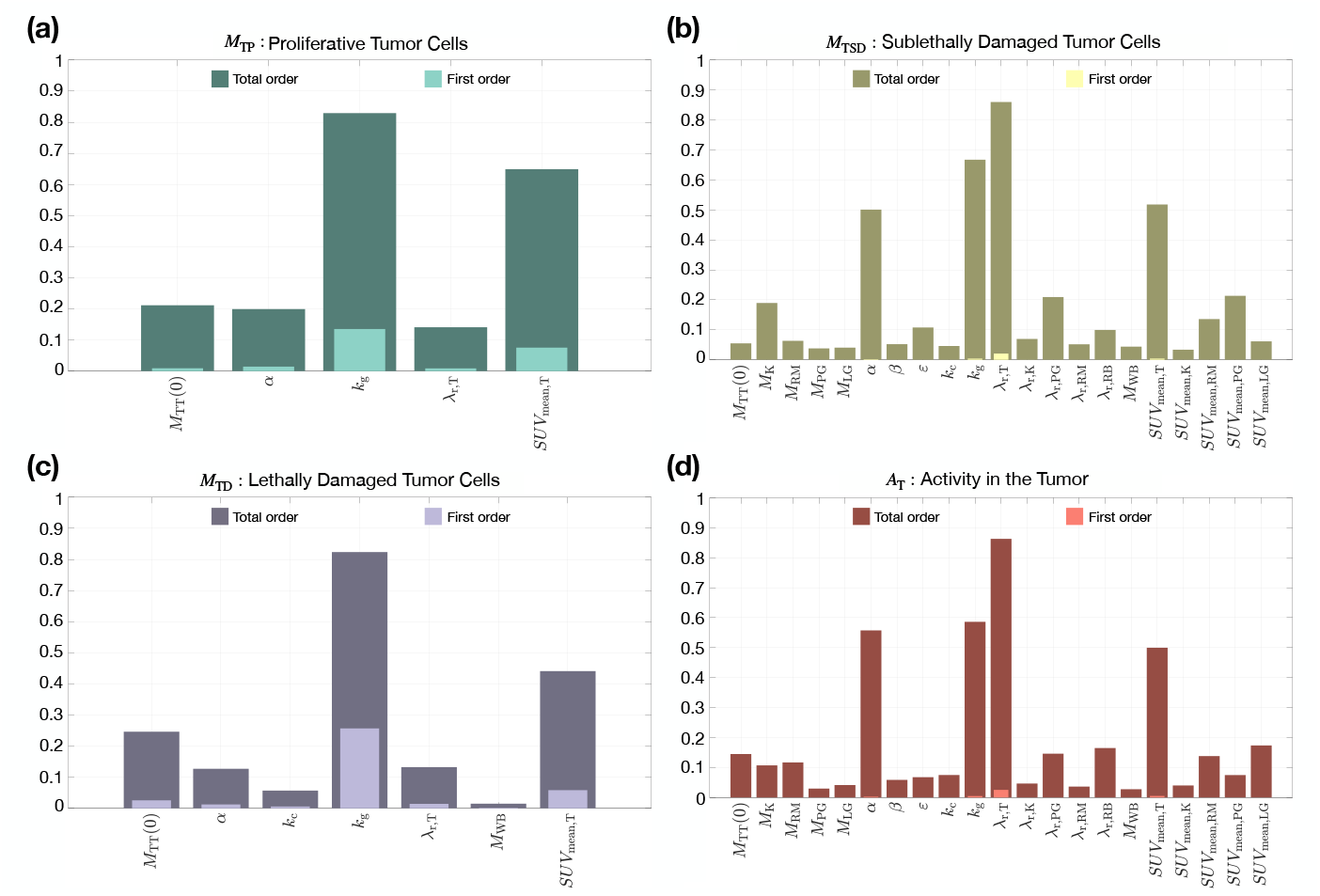
First- and total-order Sobol sensitivity indices, computed for the system (5), using polynomial chaos expansion as detailed in Section 2.6. All parameters were varied within the ranges given in Table 1. Parameters with sensitivity indices below 10^*−*2^ were omitted for clarity.

Fig. 9 shows the Sobol first- and total-order indices of the model parameters, omitting any terms with sensitivity values below a threshold of 10^−2^ for clarity. Figs S1 - S4 display the full set of indices for all parameters and populations without any threshold. Since no literature bounds are available for the sublethal-lesion interaction probability *ε*, we sampled its full feasible range (0, 1); consequently, even when *S*_1_ for *ε* is small in a given output, *ε* can still contribute through interaction effects (reflected in *S*_*T*_ ) via 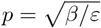 and the transition rate *k*_pp_.

Fig 9 (a), for proliferating cells (*M*_TP_), shows that the growth rate *k*_g_ dominates (*S*_*T*_ ≈ 0.83; *S*_1_ ≈ 0.13), indicating that expansion dynamics are mainly controlled by the intrinsic growth capacity of the viable tumor cell population. The mean SUV in the tumor, SUV_mean,T_ (*S*_*T*_ ≈ 0.65; *S*_1_ ≈ 0.1), also played a substantial role, reflecting the influence of tumor uptake efficiency on the AD distribution [63]. The initial proliferative mass *M*_TP_(0) and the lethal-damage coefficient *α* (both *S*_*T*_ ≈ 0.2; *S*_1_ ≈ 0.09 for *M*_TP_(0) and *S*_1_ ≈ 0.03 for *α*) contributed moderately, suggesting that tumor burden and radiosensitivity have non-negligible effects on dynamics of treatment response. The biological clearance rate *λ*_r,T_ (*S*_*T*_ ≈ 0.14; *S*_1_ ≈ 0.08) showed a moderate impact, consistent with previous findings on the influence of tracer clearance on tumor activity kinetics [64].

In Fig 9 (b), for sublethally damaged cells (*M*_TSD_), the biological clearance rate of the tumor *λ*_r,T_ (*S*_*T*_ *≈* 0.85; *S*_1_ *≈* 0.03), the tumor growth rate *k*_g_ (*S*_*T*_ *≈* 0.68; *S*_1_ *≈* 0.02), the mean SUV in the tumor, SUV_mean,T_ and the linear coefficient *α* (*S*_*T*_ ≈ 0.5; *S*_1_ ≈ 0.01), emerged as the most relevant drivers. These parameters largely determine the balance between cellular damage and repair within the tumor compartment, highlighting the importance of growth kinetics and radiation sensitivity in shaping treatment outcomes [65]. In particular, parameters such as *M*_K_, *λ*_r,PG_, and SUV_mean,PG_ (with *S*_*T*_ values approaching 0.2), as well as *ε* and *λ*_r,RB_ (with *S*_*T*_ ≈ 0.1), contribute substantially only through higher-order effects. Similarly, parameters including *M*_TT_(0), *M*_RM_, *β, k*_c_, *λ*_r,K_, *λ*_r,RM_, SUV_mean,K_, and SUV_mean,LG_ show moderate total-order sensitivity (around *S*_*T*_ *≈* 0.05) driven purely by interactions. These parameters influence *M*_TSD_ only when combined with others because their effects propagate indirectly through clearance pathways, decay of inter-compartment activity, or initial mass redistribution, none of which produce a noticeable isolated effect, but become relevant when coupled with tumor kinetics and activity-dependent damage rates.

Fig 9 (c), shows that for lethally damaged cells (*M*_TD_), *k*_g_ (*S*_*T*_ ≈ 0.82; *S*_1_ *≈* 0.25), SUV_mean,T_ (*S*_*T*_ ≈ 0.44; *S*_1_ ≈ 0.05) and *M*_TP_(0) (*S*_*T*_ ≈ 0.25; *S*_1_ ≈ 0.03) dominated. This indicates that the initial tumor proliferative state, tumor growth rate, and tracer uptake have the strongest influence on the accumulation of lethally damaged cells [66]. Contributions from *α* and *λ*_r,T_ (with *S*_*T*_ *≈* 0.13 and *S*_1_ *≈* 0.014) were moderate, while contributions from *k*_c_ and *M*_WB_ remained small (both with *S*_*T*_, *S*_1_ *<* 0.1).

Finally, in Fig 9 (d), for tumor activity (*A*_T_), *λ*_r,T_ (*S*_*T*_ *≈* 0.86; *S*_1_ *≈* 0.025), *k*_g_ (*S*_*T*_ *≈* 0.58; *S*_1_ *≈* 0.0045), *α* (*S*_*T*_ *≈* 0.55; *S*_1_ *≈* 0.024), SUV_mean,T_ (*S*_*T*_ *≈* 0.5; *S*_1_ *≈* 0.005) stood out, because growth-driven cell turnover and tracer uptake shape activity kinetics [63]. All parameters with *S*_*T*_ *<* 0.1 and negligible *S*_1_—namely *β, ε, k*_c_, *λ*_r,K_, *λ*_r,RM_, *M*_PG_, *M*_LG_, *M*_WB_, SUV_mean,K_, and SUVmean,PG—clustered together, showing only very weak effects arising purely from higher-order interactions. The remaining parameters with moderate total-order influence (0.1 *< S*_*T*_ *<* 0.2)—*M*_T_(0), *M*_K_, *M*_RM_, *λ*_r,PG_, *λ*_r,RB_, SUV_mean,RM_, and SUV_mean,LG_—affected *A*_T_ exclusively through interaction-driven pathways, mirroring the pattern observed for *M*_TSD_. Overall, these findings indicate that tumor activity is governed predominantly by tumor-specific kinetics and uptake, whereas anatomical masses and non-tumor clearance rates contribute only through higher-order interactions, reinforcing a consistent mechanistic hierarchy across model outputs.

In Supplementary Figures S5 - S8, we present the sensitivity analysis for activity in kidneys, red marrow, parotid glands, and lacrimal glands, respectively. Across the three organs kidney, red marrow, and parotid gland, the parameter with the highest sensitivity was consistently the organ-specific clearance rate *λ*_r,X_ (with *X ∈* K, RM, PG), showing *S*_*T*_ *≈* 0.98 and *S*_1_ ranging between approximately 0.70 and 0.85. For *A*_K_, in addition to the dominant contribution of *λ*_r,K_, a secondary influence arose from SUV_mean,K_ (*S*_*T*_ *≈* 0.16, *S*_1_ *≈* 0.012). For *A*_RM_, all remaining parameters contributed weakly (*S*_*T*_ *<* 0.1), although they exhibited small non-zero effects across the model. For *A*_PG_, only *M*_PG_ and SUV_mean,PG_ added minor contributions (both with *S*_*T*_ *<* 0.1), with activity still overwhelmingly governed by *λ*r,PG. In general, the predominance of *λ*_r,X_ in the kidneys, red bone marrow, and parotid glands highlights that organ-specific clearance is the central biological factor determining time-activity curves in highly renewable tissues.

In contrast, *A*_LG_ exhibited a clearly different sensitivity pattern: the dominant contributor was SUVmean,LG (*S*_*T*_ *≈* 0.40, *S*_1_ *≈* 0.39), followed by *M* WB (*S*_*T*_ *≈* 0.35, *S*_1_ ≈ 0.34) and *M*_LG_ (*S*_*T*_ ≈ 0.26, *S*_1_ ≈ 0.25), making the lacrimal gland the only compartment whose activity was not shaped by clearance-related parameters. This distinct behavior reflects the fact that, when biological clearance is fixed or negligible, activity becomes predominantly determined by tracer uptake and compartmental volume, underscoring the uptake-driven kinetics characteristic of small secretory organs.

In Fig S9, for the activity of the remaining-body (*A*_RB_), the dominant contributor was *λ*_r,RB_ (*S*_*T*_ *≈* 0.96, *S*_1_ *≈* 0.62). Secondary influences came from *k*_g_ (*S*_*T*_ *≈* 0.29, *S*_1_ ≈ 0.007) and *α* (*S*_*T*_ ≈ 0.12, *S*_1_ ≈ 0.0012). All remaining parameters showed only minor effects, with total-order indices below 0.1 and first-order indices effectively zero.

In Fig S10, for cumulative AD in tumor, *λ*_r,T_ (*S*_*T*_ *≈* 0.73; *S*_1_ *≈* 0.39), SUV_mean,T_ (*S*_*T*_ *≈* 0.41; *S*_1_ *≈* 0.13) and *M*_WB_ (*S*_*T*_ *≈* 0.04; *S*_1_ *≈* 0.0017) dominated.

In Fig S11, the sensitivity indices for the total tumor mass (the sum of *M*_TP_, *M*_TSD_, and *M*_TD_) closely mirrored the dominant drivers identified in the individual subpopulations. As expected for an aggregate quantity, the most influential parameters were the tumor growth rate *k*_g_, the tumor uptake SUV_mean,T_, the tumor biological clearance rate *λ*_r,T_, the linear radiosensitivity coefficient *α*, and the initial tumor mass *M*_T_(0). Together, these parameters govern proliferation, uptake-mediated dose deposition, radiation-induced lethality, and the initial tumor burden, thereby jointly determining the overall tumor-mass dynamics.

Finally, in Fig S12, the sensitivity indices for the whole-body activity reflected the combined influence of all organ compartments. As expected for a sum of activities, the dominant contributors were the organ-specific biological clearance rates *λ*_r,X_ (across kidneys, red marrow, parotid gland, remaining body, and tumor), together with the tumor growth rate *k*_g_, the linear radiosensitivity coefficient *α*, the lacrimal-gland uptake SUV_mean,LG_, and the tumor uptake SUV_mean,T_. These parameters collectively dictate the balance between biological clearance, proliferation-driven turnover, and tracer affinity, thereby shaping the global whole-body activity profile.

## 4 Discussion

In this study, we developed a mathematical model describing the response of mCRPC to [^177^Lu]Lu-PSMA-617. The model accounts for [^177^Lu]Lu activity in the tumor and the OARs, incorporates tumor growth dynamics and radiation-induced damage, and enables a mechanistic evaluation of how RPT scheduling influences both efficacy and toxicity in the context of mCRPC.

Previous works have addressed the mechanistic modeling of radioligand therapy in mCRPC from different perspectives. Ribes *et al*. [67] employed 3D Monte Carlo personalized dosimetry to demonstrate improved outcomes with optimized dosing. Golzaryan *et al*. [68] modeled amino-acid infusion to reduce kidney dose by enhancing renal clearance, coupling a personalized physiologically based pharmacokinetic (PBPK) model with convection–diffusion–reaction equations to account for interstitial fluid pressure and tumor heterogeneity. In four personalized cases, amino-acid infusion modified the feasible injected activity and shifted the tumor/OAR dose trade-off. Kletting *et al*. [13, 28] developed a PBPK (and later PBPK-pharmacodynamic) framework with parameter fitting to multimodal imaging. Siebinga *et al*. [69] developed a pharmacokinetic–pharmacodynamic model for [^177^Lu]Lu-PSMA-I&T that captured organ and tumor uptake, declining tumor accumulation across cycles, and prostate-specific antigen response, providing a framework to predict individual treatment outcomes and identify patients at risk of therapy failure. Hardiansyah *et al*. [70] reviewed population pharmacokinetic modeling in RPT. Finally, Piranfar *et al*. [14] analyzed spatiotemporal transport, showing minimal impact of geometry and necrotic-zone size on tumor AD.

Unlike previous studies that addressed specific aspects of RPT, such as dosimetry optimization, pharmacokinetic modeling, or localized exposure-response analysis, our work introduces an integrated, cell population-based framework that links pharmacokinetics, tumor dynamics, radiation response, and clinical outcomes simultaneously. By combining a mechanistic model with large-scale VP simulations, we were able to quantify efficacy and toxicity across a broad range of treatment schedules. This allowed us to reproduce key clinical trial results and systematically explore different regimen designs. This approach bridges the gap between mechanistic modeling and clinical translation by providing a statistically robust *in silico* platform for evaluating scheduling strategies and supporting RPT personalization.

Validating our *in silico* trials against clinical data demonstrated the model’s ability to replicate organ and tumor ADs and to predict patient survival following [^177^Lu]Lu-PSMA-617. We adopted a VP approach rather than digital twins (parameters were sampled rather than individually calibrated), so although individuals included in our virtual cohort may not match specific patients from the clinical trials, no difference would be expected at the population level. ADs from our simulations substantially overlapped with those reported by Violet *et al*. [18] and Scarpa *et al*. [19], with particularly close agreement for mean tumor dose. The largest discrepancies were observed in the lacrimal glands, likely reflecting dosimetry challenges related to small organ size and inter-patient biodistribution variability [18]. Survival validation was performed using data from two clinical studies: TheraP [20] and LuPSMA [21]. In the TheraP trial, the restricted mean survival time was reported as 19.6 months, while our *in silico* model predicted 19.47 months, with a log-rank *p*-value of 0.63, confirming statistically indistinguishable survival distributions. In the LuPSMA trial, which provided the median overall survival, our simulated value of 14.62 months closely reproduced the clinical result of 13.5 months, with a log-rank *p*-value of 0.29, also indicating no significant difference between survival curves. These findings suggest that our *in silico* approach captures the pharmacokinetics and dosimetry of [^177^Lu]Lu-PSMA-617 both in OARs and tumor lesions, while reproducing tumor-burden dynamics and their impact on survival. The reliability of the model-based predictions of survival and toxicity supports its use as a test bed to explore different therapeutic scenarios and RPT personalization. Prospective clinical trial design using these methods remains an open goal [71].

Virtual cohorts with *N* = 500 VPs provided statistically robust population-level insights. Such analysis of virtual cohort size provides an empirical framework to assess sampling reliability in population-based *in silico* studies [60]. This is often overlooked, yet it is critical for ensuring that model-driven conclusions generalize beyond a single VP sample. Such robustness is essential for informing real-world decisions, including trial feasibility, regimen design, and patient eligibility criteria.

Our sensitivity analysis revealed the impact of model parameters on survival and toxicity, with patterns consistent with clinical observations. As shown in Fig 9 (a), faster-growing clones replenish targetable cells during therapy cycles, consistent with repopulation dynamics described in radiobiology of solid tumors [72, 73]. SUV_mean,T_ governs the fraction of injected activity retained in the tumor and correlates with AD and outcomes estimated from pre-therapeutic PSMA PET, a relationship reinforced in the VISION (phase III trial of [^177^Lu]Lu-PSMA-617 in mCRPC) secondary analysis showing whole-body tumor SUV_mean_ as the best predictor of survival benefit with [^177^Lu]Lu-PSMA-617 [23]. The initial proliferative tumor mass (*M*_TP_(0)) and lethal-damage coefficient (*α*) encode tumor burden and radiosensitivity within the linear-quadratic framework [64], while tumor biological clearance (*λ*_r,T_) modulates intratumoral retention [74].

The results for sublethally damaged cells (*M*_TSD_) shown in Fig 9 (b), are biologically consistent: the balance between activity-mediated lesion induction (driven by tumor uptake and residence time), proliferative replenishment of surviving cells, and intratumoral clearance underpins the dynamics of sublethal damage: faster intratumoral clearance shortens residence time and reduces ongoing lesion induction, whereas tumor growth (repopulation) and intrinsic radiosensitivity (*α*) govern the flux of cells between damaged and viable compartments [75, 76, 77, 78]. Notably, parameters regulating the progression out of the sublethal compartment (e.g., the interaction probability *ε* entering into *k*_pp_) exhibited low total-order indices but essentially zero first-order terms, indicating that their influence emerges only through interactions rather than direct individual effects. Comparable interaction-driven behavior is reported in mechanistic and PBPK models, where receptor binding, perfusion and clearance jointly shape residence times and dose accumulation [79, 80]. Proliferative replenishment and tracer uptake dominate lethally damaged cells [66], as shown in Fig 9 (c).

The growth- and uptake-driven kinetics are fundamental to tumor activity (*A*_T_)[63], as shown in Fig 9 (d). For normal tissues (Figs S5-S7), the analysis consistently shows that organ-specific biological clearance rates (*λ*_r,X_, where X ∈ {K, RM, PG}) are the dominant determinants of organ activity and hence organ dose for kidneys, red marrow, and parotid glands; uptake (organ SUV_mean_) and organ mass play secondary roles. This behavior reflects well-known physiology (differential perfusion, excretory function and PSMA-ligand handling produce marked inter-organ differences in residence time that control cumulative exposure and toxicity risk) and supports dosimetry-guided strategies that prioritize measuring or constraining organ clearance to mitigate toxicity (e.g., renal protection strategies and salivary-sparing approaches) [81, 41].

For the activity of the lacrimal gland, *A*_LG_ (Figs S8), the clearance parameter was fixed and thus excluded from the sensitivity analysis; accordingly, the most influential parameters were SUV and mass. Biologically, this means that when local elimination is not variable, the absorbed dose is primarily determined by how much radioligand is taken up and how large the gland is; small secretory organs with high affinity for the ligand can accumulate a large dose per unit activity, even if their absolute uptake is modest. This finding is in line with dosimetry results in prostate cancer RPT: for example, a meta-analysis of [^177^Lu]Lu-PSMA therapies showed that the lacrimal glands appear among the sites with the highest absorbed doses per GBq, frequently exceeding that of salivary glands [25]. Such dose accumulation, independent of clearance variation, underlines the potential risk for toxicity in highly affine, small-volume organs and supports strategies for local dose mitigation when uptake is elevated (e.g., cooling or optimized fractionation) [41].

Fig S9 reflects the fact that whole-body activity depends not only on how rapidly the unbound radioligand is eliminated but also on how quickly the tumor compartment expands and demands tracer redistribution. Faster proliferating tumors behave as larger biological sinks, transiently reducing circulating activity and consequently altering the kinetics of systemic retention. Biologically, this interplay is consistent with observations in PSMA-targeted therapies, where tumor burden and proliferation modulate whole-body pharmacokinetics by sequestering variable fractions of circulating radioligand [82, 18]. Thus, variability in both systemic clearance and tumor growth jointly shapes global activity dynamics, highlighting the importance of patient-specific tumor kinetics when estimating whole-body exposure and potential toxicity.

The hierarchy of Fig S10–cumulated absorbed dose in the tumor–reflects the fundamental relationship between retained tumor activity and delivered dose: longer intratumoral residence and higher ligand affinity directly increase energy deposition, which is consistent with clinical evidence. This shows that tumor SUV_mean_ and effective half-life are the strongest predictors of absorbed dose and response in PSMA-targeted radioligand therapy [63, 18]. Biologically, these findings reinforce that therapeutic efficacy in PSMA-RPT is governed not by instantaneous uptake alone but by the interplay between affinity-driven internalization and slow efflux kinetics, both of which dictate how much of the administered activity is ultimately converted into a cytotoxic dose.

From the simulated treatment regimens, several insights emerged. First, intensive regimens better limit inter-cycle regrowth: fewer, higher-activity injections slightly improved mOS, with the one-injection and asymmetric two-injection schedules yielding the longest mOS values. However, these survival gains were balanced by increased toxicity, illustrating the central efficacy–safety trade-off inherent in RPT. Asymmetric schedules offered a promising compromise, enhancing survival with only a modest rise in renal and parotid gland toxicity. Conversely, delivering the same standard fixed total activity (44.4 GBq [6]) in a larger number of injections beyond nine yielded progressively diminishing mOS, suggesting that concentrating dose temporally can improve efficacy in a subset of patients.

We also explored cycle timing under six-injection (7.4 GBq each) protocols [6]. Our results showed that mOS and toxicity were relatively insensitive to cycle length within a broad window (up to 9 weeks), offering flexibility for clinical logistics and constraints. However, excessively delayed cycles (beyond 12 weeks) resulted in sharp declines in survival, likely due to tumor regrowth or loss of treatment continuity due to patient death. Interestingly, these longer cycles were associated with reduced organ toxicities, suggesting that moderately delayed schedules could be feasible for patients at higher risk of toxicity, albeit at the cost of some loss in efficacy. Indeed, we showed that patients could substantially reduce their toxicity risk with a more spaced protocol (cycle length of 9 weeks), while achieving survival outcomes comparable to the standard 6 week regimen–more intensive and associated with higher toxicity risk.

Negligible lacrimal and small parotid gland toxicities were observed across simulated schedules, suggesting these organs are rarely dose-limiting for mCRPC under [^177^Lu]Lu-PSMA, whereas kidneys and red marrow emerged as the primary dose-limiting OARs, in line with clinical experience [83, 84, 85].

Our results are generally consistent with those of Zaid *et al*. [17], who identified the 2* protocol as a favorable administration strategy, improving OS while maintaining acceptable toxicity in OARs. However, our findings indicate an improvement in survival (around 17% with respect to the standard regimen) accompanied by a modest increase in the probability of toxicity in kidneys (PoT_K_), red marrow (PoT_RM_), and parotid glands (PoT_PG_) under the 2* protocol, specifically +14.4%, +3% and +7.2%, respectively, with respect to the standard protocol [6]. This suggests that, while some patients (around 30% of VPs) may benefit substantially from this regime, others may be at higher risk of developing toxicity (around 15% of VPs). Note that around 50%, 30%, and 10% of VPs already faced kidneys, red marrow, and parotid glands toxicological issues, respectively, under standard treatment, in line with clinical trial observations. Indeed, Santo *et al*. [86] reported long-term toxicities in 8 of 18 (44%) patients, and serious toxicities occurred in 3 of 8 patients (specifically one thrombocytopenia, i.e., red marrow toxicity, and two renal failures, i.e., kidneys toxicity). Hermann *et al*. [83] observed marrow suppression in 36.7% of patients. Mahajan *et al*. [85] reviewed salivary gland toxicity in several studies, reporting results consistent with our simulations. For instance, Rahbar *et al*. [87] observed xerostomia (salivary/parotid glands toxicity) in 4 of 28 (14.3%) patients and it was not permanent. These observations underscore the importance of inter-individual variability when evaluating protocols.

Overall, these results delineate a large therapeutic window for [^177^Lu]Lu-PSMA RPT. Cycle lengths between 2 and 9 weeks provide comparable survival and toxicity outcomes, affording clinicians flexibility in scheduling. Among dose-fractionation options, the 2* regimen provided a slighlty higher mOS (close to the single-injection schedule) while mitigating kidney and parotid toxicity. The standard 6 × 7.4 GBq protocol achieved a smaller mOS but led also to less toxicity, remaining a robust and practical balance between efficacy, safety, and logistical convenience. For more conservative scenarios prioritizing toxicity reduction, the eight-injection schedule provided a reasonable compromise. These insights suggest that the standard protocol is a good option at the population level. However, some patients could achieve improved survival without experiencing toxicity with different treatment strategies, paving the way for personalized medicine.

Our framework has several strengths. It enables efficient simulation of large virtual patient cohorts and supports robust statistical comparisons (including pairwise survival analyses). The model integrates pharmacokinetics, tumor growth, and radiation response while remaining computationally tractable [88]. Parameters can be estimated from patient-specific data, enabling digital-twin applications; importantly, several of the most influential parameters could be derived from images (see e.g. [89]). We incorporated convergence testing to ensure population-level reproducibility and trustworthy clinical translation [60], features that promote both scientific rigor and clinical utility.

From the broader perspective of RPT or TRT, the molecular blueprint outlined by Primac et al. [2] emphasizes several open challenges for TRT optimization, including the need to balance tumor retention against off-target exposure in kidneys, bone marrow, and salivary glands, to implement individualized dosimetry in routine practice, to exploit quantitative molecular imaging for patient selection and response assessment, and to rationally optimize treatment schedules and radioligand properties within a narrow therapeutic window. Our population-based framework directly addresses these questions at the macroscopic level by linking imaging-derived uptake and clearance parameters to organ-specific dose constraints and survival outcomes, thereby quantifying how interpatient variability in pharmacokinetics and organ tolerance shapes the efficacy-toxicity trade-off of a given regimen. In the specific case of [^177^Lu]Lu-PSMA-617, our approach delineates a therapeutic window for cycle timing (2–9 weeks) and identifies regimens that offer favorable population-level efficacy–toxicity balance under fixed total activity. Moreover, the dominant influence of tumor SUV_mean_ and initial tumor burden on simulated survival in our global sensitivity analysis provides a mechanistic underpinning for the blueprint’s emphasis on molecular imaging–derived metrics as key tools for patient stratification and trial enrichment. Furthermore, because pharmacokinetics and radiobiology are encoded in a modular way, our model can be readily adapted, via reparameterization of uptake and clearance, to emulate next-generation ligands or radioprotective strategies proposed in the TRT roadmap without redesigning the overall framework. For example, hypothetical compounds could be explored with altered clearance or improved tumor retention, and schedules incorporating nephroprotective amino-acid infusions be evaluated to anticipate their impact on tumor control and organ doses before clinical testing. In this sense, our *in silico* platform provides a quantitative companion to experimental and clinical efforts aimed at realizing the individualized, dosimetry-guided TRT paradigm envisioned in the TRT molecular blueprint by Primac et al. [2].

Finally, our model can be adapted to other tumors and radioligands, such as neuroendocrine tumors under [^177^Lu]Lu-DOTATATE [7] or thyroid cancer under ^131^I-NaI [90]. Updating organ kinetics and radiobiology would also allow evaluation of alternative isotopes/targets (e.g., SSTR2). The model’s modular structure facilitates such extensions, supporting a broader role for *in silico* trials in RPT optimization.

This study has several limitations. First, VP generation used ranges from specific studies and assumed uniform parameter distributions, which may not fully capture real-patient variability [91]. Second, toxicity models do not describe different grades of toxicity, assume uniform organ responses, and do not capture inter-individual radiosensitivity. Third, the model lacks spatial resolution, which may matter for size- and morphology-dependent efficacy [92]. Finally, we focused on systematic exploration rather than formal optimization, and clinical validation was limited to retrospective comparisons with published trials.

Future work could address these limitations and incorporate patient-specific imaging together with physiological data to personalize VP profiles (digital twins), improve predictive accuracy [55] as well as treatment outcomes. Indeed, our results suggest that treatment should be personalized to each patient since no schedule emerged clearly as optimal at the population level. Specifically, patient-specific image data could be used to calibrate the OAR and tumor SUV parameters, genomic predictors for radiosensitivity, and prostate-specific antigen levels for tumor growth rate [22]. Such a personalized model could be used to tailor treatments to individuals. Moreover, personalized *in silico* trials could inform trial design, patient selection, or real-time treatment adaptation, bringing computational modeling closer to the clinical bedside. This positions our model as a bridge between mechanistic radiopharmacology and clinical personalized decision-making. Expanding the model to consider combination therapies or tumor resistance mechanisms could further increase clinical relevance.

Personalized frameworks based on mechanistic mathematical models and *in silico* trials offer powerful tools for preclinical assessment of complex treatment strategies such as RPT for mCRPC. Our findings highlight the importance of dose scheduling and cohort-level validation in maximizing both efficacy and reliability. As virtual populations become more sophisticated, they will increasingly complement and inform real-world clinical decision-making.

## 5 Conclusion

We developed a patient-personalizable mechanistic model of [^177^Lu]Lu-PSMA-617 therapy for mCRPC that links pharmacokinetics, organ dosimetry, radiobiology, and tumor-mass dynamics, enabling robust *in silico* trials. The framework reproduces organ and tumor absorbed doses and survival outcomes reported in clinical studies and, using an empirically validated virtual cohort of 500 virtual patients, provides a reliable platform to test risk-aware treatment strategies.

Across fixed total activity, fewer, higher-activity injections modestly improved median overall survival but increased particularly renal and parotid toxicity; asymmetric dosing (e.g., two-injection schedules with unequal activities) offered a favorable efficacy–toxicity trade-off. Cycle timing was flexible within a broad window (2-9 weeks) with negligible impact on survival but spaced regimens reduced toxicity, whereas delays greater than 12 weeks reduced survival. No single schedule emerged as universally optimal, underscoring the need for personalization-guided by imaging-derived uptake and clearance parameters-to maximize patient benefit while respecting organ-at-risk constraints. Our results support the use of the modeling approach proposed here as a quantitative tool to design and personalize risk-aware RPT protocols.

## Code availability

The source code used to simulate the model presented in this manuscript is available on a public GitHub repository: https://github.com/matteo-italia94/radiopharmaecutical-therapy-for-metastatic-prostate-cancer.git.

## Acknowledgments

This work has been partially supported by Ministerio de Ciencia e Innovación/Agencia Estatal de Investigación, Spain (doi:10.13039/501100011033, grant number PID2022-142341OB-I00) and European Regional Development Fund (ERDF A way of making Europe), by Junta de Comunidades de Castilla-La Mancha and European Regional Development Fund (grant numbers SBPLY/23/180225/000041 and SBPLY/24/180225/000193), and by the University of Castilla-La Mancha/ERDF, Applied Research Projects within the UCLM research programme (grant number 2025-GRIN-38309). S. B-V. has been supported by grant number 2022-GRIN-34405 funded by the University of Castilla-La Mancha/FEDER, Applied Science Projects within the UCLM research programme.

